# Sex and reproductive condition shape thermal acclimation strategy in a plethodontid salamander

**DOI:** 10.1101/2025.09.28.679080

**Authors:** Kevin J. Moore, Brittany M. Winter, Jeanette B. Moss

## Abstract

The question of whether males and females differ in their responses to temperature is essential to understanding adaptive capacities in a warming world. However, sex has traditionally been neglected in the field of thermal ecology, and our understanding of the factors that promote sex differences in thermal plasticity is underdeveloped. Here, we investigate the independent and interactive effects of sex and reproductive condition on thermal acclimation capacity in a plethodontid salamander (*Plethodon cinereus*). We carried out two stop-flow respirometry experiments, with salamanders acclimated to one of two thermal environments designed to simulate a ‘Cool’ or ‘Warm’ breeding season. In the first experiment, we compared the thermal acclimation responses of males and gravid females across a gradient of three ecologically relevant test temperatures and observed distinct patterns of sexual dimorphism of standardized metabolic rate (SMR) between thermal treatments that were not attributable to differences in body size. Specifically, gravid females in the warm treatment showed reduced SMR relative to gravid females in the cool treatment, particularly at higher test temperatures. In the second experiment, we expanded on our initial findings by directly testing the contribution of reproductive condition to observed sex differences in thermal acclimation capacity. By repeating the experiment of the early breeding season but including a third group of non-gravid females, we: (1) recapitulated our original finding – that gravid females exhibit a pronounced response to thermal acclimation relative to males; and (2) showed that female thermal acclimation responses are absent in nongravid females, and therefore we concluded that responses are contingent on reproductive condition. Taken together, our results provide a first glimpse into how sex and reproductive condition contribute to intraspecific variation in thermal acclimation capacity in an amphibian and underscore the need for more hypothesis-driven studies to directly test when, where, and how such patterns arise.

## Introduction

The capacity of an organism to acclimate its biological rates in response to changing thermal conditions (i.e., thermal acclimation capacity) is a widespread, often beneficial form of phenotypic plasticity that increases individual tolerance and performance at a given temperature and may contribute to population resilience under rapidly changing climates (Angilletta 2009; Rohr et al. 2018; Havird et al. 2020). Strategies such as the active up-or down-regulation of metabolism relative to basal rates can be an effective mechanism for balancing energy losses and gains at temperatures that are suboptimal for activity (Angilletta 2009; Riddell, Odom, et al. 2018; Havird et al. 2020). For example, heat-induced reductions in metabolic rate (i.e., *metabolic depression*) are common in ectotherms living in warm environments and presumably magnify energy savings during extended bouts of inactivity (Feder et al. 1982; Christian et al. 1999; Guppy and Withers 1999; Liao et al. 2021; Muñoz et al. 2022), whereas mitigating prohibitively cold temperatures often involves inflating metabolic rates above expected levels to extend the range of physiological function (i.e., *metabolic compensation*; (Rogers et al. 2007; Gaston et al. 2009; Bozinovic et al. 2011; Catenazzi 2016; Watanabe and Payne 2023)). Accounting for this plasticity can dramatically alter ecological predictions under climate change, and is therefore imperative to accurately modeling population responses (Riddell, Odom, et al. 2018; Briscoe et al. 2023). However, species- and population-level measures of thermal acclimation capacity are often derived from few reference individuals, which may not accurately reflect the population as a whole (Bennett et al. 2019). Indeed, recent work has documented tremendous inter-individual variation in thermal acclimation strategies (Seebacher et al. 2015; Riddell, McPhail, et al. 2018; Hui et al. 2020; Liao et al. 2021), although knowledge of the factors that account for this variation remains fragmentary (Bennett et al. 2019; Bodensteiner et al. 2021; Cocciardi and Ohmer 2024).

One poorly understood axis of inter-individual variation in thermal acclimation capacity is sex (Pottier et al. 2021; Hangartner et al. 2022). Male and female animals differ in a broad range of morphological, physiological, and behavioral traits. These adaptations are often exaggerated during reproductive periods, when the sexes tend to conform to stereotyped patterns of energy storage and expenditure associated with their distinct reproductive strategies. Where females typically adopt conservative strategies, accumulating and provisioning energy gradually towards costly egg production, males tend to take more risks, accumulating energy in short bursts and expending reserves in active pursuit of reproductive opportunities (Beck et al. 2003; Czenze et al. 2017; Tarka et al. 2018). Such distinct approaches to energy use may favor different strategies for coping with thermal variation. For example, a study of the striped marsh frog uncovered male-biased metabolic compensation responses to cold temperatures during the early spring breeding season (Rogers et al. 2007), suggesting that thermal acclimation may be enhanced in males relative to females when there is a benefit to mating performance. Conversely, several studies have revealed female bias in capacity to acclimate to high temperatures, which could reflect their slow pace-of-life and high reproductive investment (Edmands 2021; Pottier et al. 2021; Vermandele et al. 2024). Because population growth is often more sensitive to the survival and fecundity of one sex (e.g., ‘female demographic dominance’), the potential for sex-biased plasticity presents a critical, albeit relatively unexplored factor influencing population persistence (Fox et al. 2019; Hangartner et al. 2022; Gissi et al. 2023).

Despite growing interest of recent years (e.g., (Huey and Pianka 2007; Lailvaux 2007; Bodensteiner et al. 2021; Edmands 2021; Gunderson 2021; Pottier et al. 2021; Hangartner et al. 2022)), the number of empirical studies explicitly testing for sex differences in thermal response continues to lag behind theory. In their 2021 meta-analysis, Pottier et al. found that over three quarters of eligible studies on thermal acclimation capacity (446 of 586) failed to report or did not consider the sex of experimental subjects. Of the remaining eligible studies, overall effects were heterogeneous and supported few systematic patterns, suggesting that where it arises, sexual dimorphism in thermal response may be context dependent. For instance, countless field studies have documented variation in thermal traits associated with reproductive condition or breeding stage (e.g., (Charland and Gregory 1990; Grinevitch et al. 1995; Finkler et al. 2003; Galliard et al. 2003; Hammill et al. 2004; Isaac and Gregory 2004; Finkler 2006; Rogers et al. 2007; Wilson et al. 2007; Darnell et al. 2013; Allen and Levinton 2014; Beal et al. 2014; Vaughn et al. 2014; Webber et al. 2015; Woolrich-Piña et al. 2015; Parker et al. 2019; Levine et al. 2024), which could confound detection of sex differences. Given practical limitations of acquiring detailed physiological and behavioral data from multiple demographic classes when forecasting climate change responses, there is a need to identify when, where, and how sex-specific responses to temperature are likely to emerge in nature (Pottier et al. 2021; Hangartner et al. 2022).

Lungless salamanders (Family: Plethodontidae) are well suited for investigating the role of sex and reproductive condition in driving thermal acclimation patterns. As small-bodied amphibians with small home ranges and narrow climatic tolerances, plethodontids rely on physiological acclimation mechanisms to function successfully across their natural geographic range (Gifford and Kozak 2012; Markle and Kozak 2018; Riddell et al. 2024), as well as in the face of climatic extremes (Novarro et al. 2018; Riddell, Odom, et al. 2018; Muñoz et al. 2022). Yet the potential for sex-specific selection pressures to shape patterns of thermal acclimation has received almost no attention in climate projections for this group. Addressing this limitation is critical, as reproduction profoundly impacts energy balance (Fitzpatrick 1973; Crespi 2001; Finkler and Cullum 2002; Takahashi 2002; Gillette 2003) and behavior (Gergits and Jaeger 1990; Dyal 2006; Leclair et al. 2008; Jaeger et al. 2016) and in many species spans the entire year (Crespi 2001; Fisher-Reid et al. 2024). Moreover, reproductive life history strategies tend to be highly sexually dimorphic in plethodontids. For example, whereas male gamete production is a relatively rapid process that relies minimally on stored energy and occurs annually (Sayler 1966; Werner 1969; Nagel 1977; Takahashi and Pauley 2010; Fisher-Reid et al. 2024), females spend up to nine months yolking a single clutch of eggs, and may reproduce as infrequently as every four years (Gillette 2003; Leclair et al. 2008; Takahashi and Pauley 2010).

Here, we enlist the eastern red-backed salamander (*Plethodon cinereus*), a species widespread across Eastern U.S. deciduous forests, to compare physiological thermal acclimation strategies between the sexes across their nearly eight-month breeding season. Due to sex differences in energy allocation to reproduction, we predicted that metabolic compensation in response to cool temperature acclimation would be male-biased, whereas metabolic depression in response to warm temperature acclimation would be female-biased. Our results provide a first glimpse into how sex and reproductive condition contribute to intraspecific variation in thermal acclimation capacity in an amphibian.

## Methods

### Study System

Mountain Lake Biological Station (MLBS) sits atop Salt Pond Mountain, a 1,160m summit located in southwest Virginia. This site has long served as the epicenter of field research on the eastern red-backed salamander and details of natural history, behavior, and breeding phenology are particularly well documented (Fisher-Reid et al. 2024). Though many populations of red-backed salamanders show color polymorphism, the “leadback” phase is virtually absent at MLBS, allowing us to focus our analyses to the level of sex. Most females at MLBS follow a biennial cycle of reproduction, such that sexually mature females fall into two equally occurring reproductive classes: gravid and non-gravid (Gillette 2003, Moss et al., unpublished). This pattern of biennial female reproduction appears to hold generally across the range of P. cinereus, and reflects the steep energetic requirements of producing direct-developing eggs given limited growing seasons to support fat accumulation (Leclair et al. 2008; Petranka 2010; Takahashi and Pauley 2010; Fisher-Reid et al. 2024). As far as we know, sexually mature male *P. cinereus* breed every year (Fisher-Reid et al. 2024). The breeding season at MLBS lasts from September–April, with the majority of mating (i.e., as evidenced from tracking occurrences of spermatophores in female vents) appearing to fall within two seasonal peaks: October and March (Gillette 2003).

Temperatures during the prolonged breeding season are highly variable but generally cool (Fig 1A). For our thermal acclimation treatments, we chose two ranges of temperatures to simulate ‘Cool,’ or current (8^°^C nights, 14.5^°^C days) and ‘Warm,’ or projected future (13^°^C nights, 19.5^°^C days) breeding season conditions. Temperatures were programmed to ramp on a 12:12 D:L photoperiod, with light and temperature cycles reversed to ensure physiological measurements aligned with the active period for nocturnal salamanders. This design would allow us to test whether salamanders acclimated to cool versus warm conditions would exhibit relatively higher metabolic rates at low temperatures or lower metabolic rates at high temperatures (metabolic depression), respectively (Fig. 1B). determine whether thermal responses vary predictably on the basis of sex and/or reproductive condition following our predictions (Fig. 1C), we assayed subjects of both sexes and reproductive classes (gravid and non-gravid).

**Figure 1.**
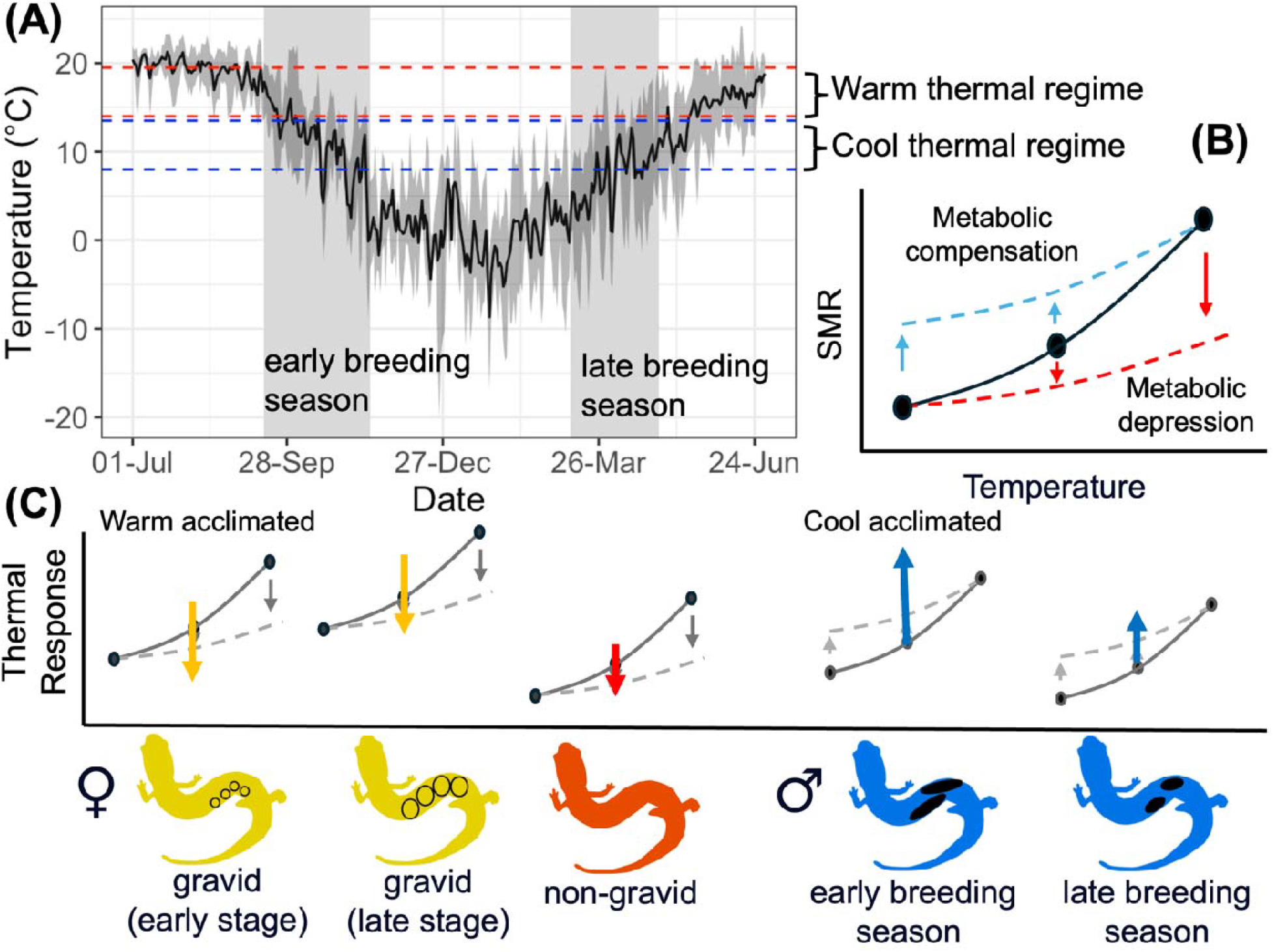
Schematic overview of thermal acclimation experiment and predictions. (A) Trends in mean ambient temperature at Mountain Lake Biological Station from 2019 to 2022. Shaded bars represent the periods of peak breeding activity in the fall (early breeding season) and spring (late breeding season). Dashed lines represent the temperature ranges chosen for the Cool thermal regime (in blue) and Warm thermal regime (in red). (B) Two predicted patterns of physiological response (depicted as standard metabolic rate, SMR) to thermal acclimation. For a given temperature-SMR relationship (black line), SMR should increase at low temperatures in response to cool acclimation (i.e., metabolic compensation, blue line) and decrease at high temperatures in response to warm acclimation (i.e., metabolic depression, red lines). (C) Predicted patterns of physiological rate and thermal response by sex and reproductive class, with black and white graphs depicting the expected variation in SMR intercepts among groups and arrows depicting the magnitude (depicted as increasing arrow length) and direction of change in SMR in response to either warm (females) or cool (males) thermal acclimation.

### Experiment 1: Early Breeding Season

The goal of our first experiment was to compare thermal responses of male and female *P. cinereus* in reproductive condition to determine whether thermal acclimation capacity represents a dimension of sex-specific plasticity in this species. We predicted that salamanders acclimated to warm thermal regimes would show depressed metabolic rates at higher test temperatures, as has been demonstrated elsewhere (e.g., (Feder 1985; Bernardo and Spotila 2006; Markle and Kozak 2018; Muñoz et al. 2022); Fig. 1B). However, if depression of metabolic rate incurs opportunity costs during the breeding season (i.e., due to insufficient delivery of oxygen for energy-demanding activities, such as mate searching), we predicted that males would opt to maintain temperature-specific metabolic rates or even increase metabolic rates when acclimated to warm breeding season conditions.

## Methods

### i. Collection and Husbandry

A total of 40 salamanders (20 females and 20 males) were collected from the vicinity of MLBS between 30 August 2024 and 20 September 2024. All females were gravid, as confirmed by candling of egg follicles (Gillette and Peterson 2001), and all salamanders had complete tails. Upon collection, we recorded the body mass (within the nearest 0.001 g), snout-vent length (SVL; distance in mm from the tip of the snout to the top of their vent), and total length (TL; distance in mm the tip of the snout to the tips of the tail) of each experimental subject. Prior to weighing, salamanders were gently patted dry with a kimwipe and induced to urinate via gently pressure on the pelvis to remove excess water weight and ensure accurate measurement. All experimental subjects were considered sexually mature adults on the basis of body size (SVL > 34mm).

Following initial processing, salamanders were distributed evenly across the two thermal acclimation treatments with respect to size and sex. Thermal regimes and photoperiods were maintained by environmental chambers (Caron Scientific 7000 series) programmed to ramp between the low temperature setpoint between the hours of 0700–1900 (reversed ‘night’) and the high temperature setpoint between the hours of 1900–0700 (reversed ‘day’). Salamanders were housed individually in 120mm petri dishes set on incubator shelves and rotated daily. A moistened paper towel, hand-misted daily with spring water, provided a saturated local humidity environment. Salamanders were fed three times per week, alternating between flightless fruit flies (*Drosophila melanogaster*) dusted with vitamins and white worms (*Enchytraeus albidus*). Paper towels were replaced once weekly.

Following recommended best practices for studying thermal acclimation in plethodontids (Homyack et al. 2010; Markle and Kozak 2018), salamanders were allowed to acclimate to their respective thermal treatments for a minimum of three weeks (range=24–38 days) before starting physiological trials. In the 10 days leading up to individual trials, food was withheld to ensure subjects were in a post-absorptive state for measuring metabolic rates (Homyack et al. 2010).

### ii. Stop-Flow Respirometry Protocol

Standard metabolic rates (SMR) were assayed across a gradient of three ecologically relevant test temperatures (8, 12, and 16^°^C) by way of automated stop-flow respirometry at a flow rate of 80 ml/min (Appendix S1). All trials were conducted between 0800 and 1600 during the day. To initiate physiological trials, salamanders (N=7 per trial) were weighed within 0.001 g, as above, before being loaded into test chambers. One chamber in the 8-channel multiplexer was left empty to collect baseline data. Three measures of volumetric O2 consumption (VO2) were recorded for each individual per trial using Expedata software, and data transformations were carried out following established manual bolus integration calculations (Lighton 2019).

Repeat testing of individual subjects occurred over five days, consisting of three days of trials alternating with one day of rest. The order of test temperatures was randomized for each set of N=7 salamanders. At the conclusion of trials, normal feeding resumed. Two repeat trials could not be completed due to tail loss (N=1 warm-acclimated male at 12^°^C and N=1 cool-acclimated female at 8^°^C). At the end of the experiment, all test subjects were euthanized and dissected to confirm sex and reproductive condition.

### iii. Statistical Analyses

All statistical analyses were carried out using R v 4.2 (R Core Development Team 2020). To analyze variation in SMR across sexes, treatments and test temperatures, we ran a series of linear mixed effects models using the package lme4 (Bates et al. 2015). For each model, log (SMR) was specified as the response variable and subject ID and trial date were specified as random effects. We first investigated whether the scaling relationship of SMR with body mass differed between the sexes or across treatments by modeling SMR for each group separately and deriving mass-scaling coefficients from the slope of the fixed effect of body mass at the start of the trial. We then ran a fully parameterized model including two-way and three-way interactions between body mass, treatment, and sex. Based on a lack of significant interaction terms, we retained body mass as a covariate but excluded statistical interactions in all subsequent models. Differences in SMR between sexes and treatments were visualized using mass-adjusted estimates, which were derived by dividing VO2 in ul/hr by starting body mass.

To test for significant thermal acclimation responses to treatment and determine whether these differed between the sexes, we expanded our base model to include sex, treatment, and their interaction as fixed effects. To investigate more specifically how males and females responded to thermal acclimation across the gradient of test temperatures, separate models were constructed for each sex including treatment, test temperature, and their interaction as fixed effects. To identify significant pairwise differences in SMR across temperatures and treatments, post-hoc tests were carried out using the package emmeans (Lenth et al. 2022).

## Results

Subjects showed marginally significant sex differences in mass upon collection (F = 3.293, *P* = 0.077), with gravid females being larger on average (1.288 ± 0.235) than males (1.148 ± 0.258). However, the initial mass of subjects did not differ between the acclimation treatments (Males: t 0.582, *P* = 0.568; Females: t = 0.666; *P* = 0.513). Mass-scaling coefficients of SMR did not differ significantly between sexes or across treatments (X^2^ = 0.008, *P* = 0.928; Fig. S1). Rather, we observed distinct patterns of sexual dimorphism of SMR between thermal treatments not attributed to differences in body size (Fig. 2). Whereas sex had no overall effect on SMR across the three test temperatures (X^2^ = 0.064, *P* = 0.801), both thermal treatment (X^2^ = 7.373, *P* = 0.007) and the interaction between sex and thermal treatment (X^2^ = 4.448, *P* = 0.035) were significant predictors of SMR. SMR generally increased with test temperature, and thermal acclimation treatment had no effect on this relationship in males (X^2^ = 1.497, *P* = 0.473). Yet a distinct pattern emerged in gravid females, who showed a strong interactive effect of thermal treatment and test temperature on SMR (X^2^ = 14.214, *P* = 0.0008). Specifically, gravid females reduced SMR in the warm treatment relative to the cool treatment (X^2^ = 10.11, P = 0.014), a trend primarily attributed to metabolic depression at the highest test temperature (16^°^C: Tukey ratio = 3.81, *P* = 0.006).

**Figure 2.**
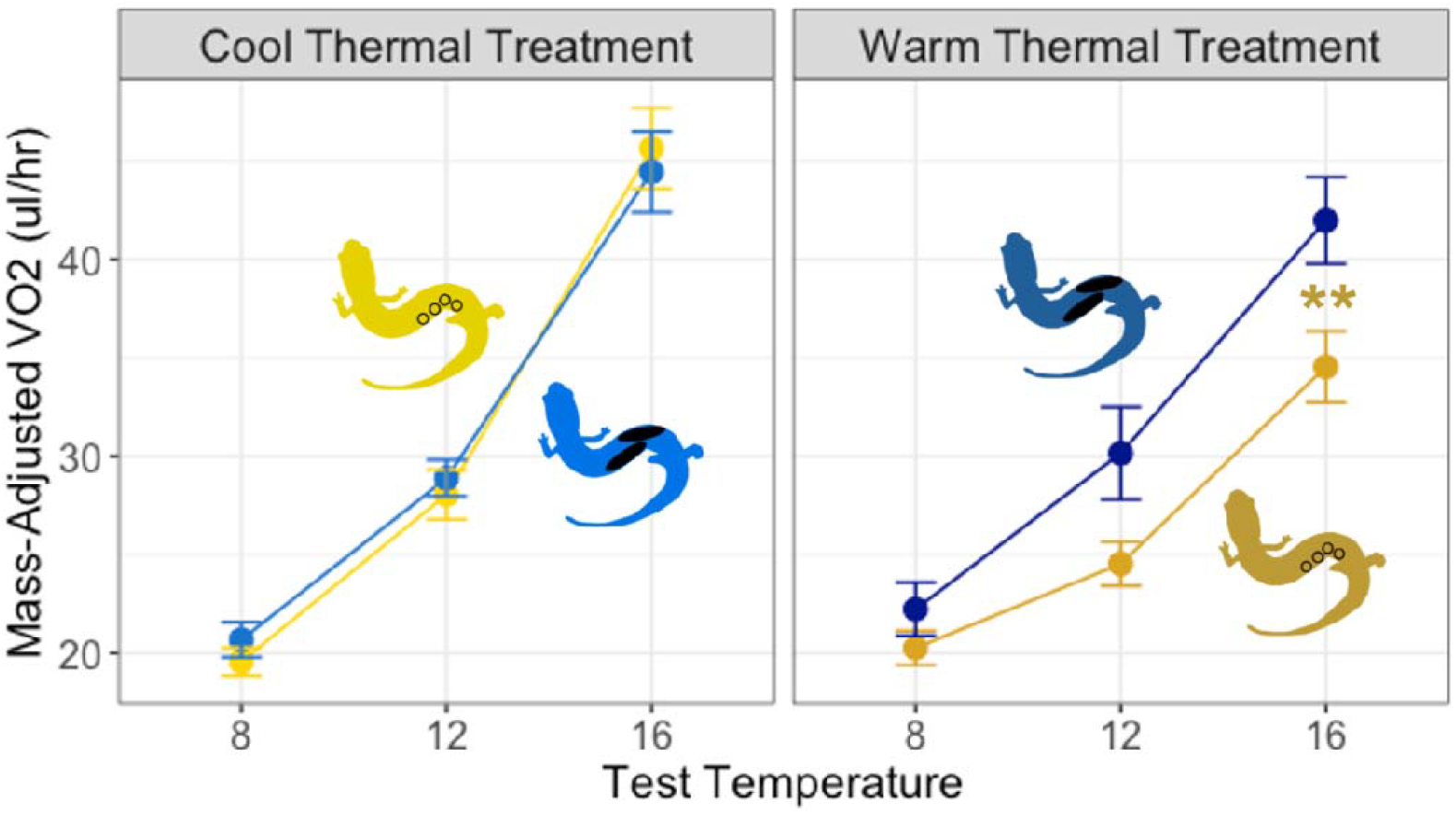
Standard metabolic rate (mass-adjusted VO2, in ul/hr) measured at three test temperatures for two classes of salamanders following 3–5 weeks thermal acclimation in one of two treatments (Cool or Warm) in the fall: gravid females (N=10 cool, N=10 warm, in gold) and males (N=10 cool, N=10 warm, in blue). Asterisks denote significant within-sex treatment differences at a given test temperature in a linear mixed effects model (* < 0.05, ** < 0.001, * * * < 0.0001).

### Experiment 2: Late Breeding Season

Building on the results of Experiment 1, the goal of our second experiment was to determine whether sex-specific patterns of thermal response in *P. cinereus* are attributable to intrinsic sex differences or to reproductive condition. A secondary goal was to track potential variation in thermal response of reproductive males and females across the reproductive season. We predicted that if sex differences in thermal plasticity are due to differences in reproductive strategy, then gravid females collected late in the breeding season should show thermal responses similar to or more pronounced than gravid females collected early in the breeding season, whereas non-gravid females would show thermal responses more similar to males (Fig. 1C).

## Methods

### i. Collection and Husbandry

A total of 84 salamanders (28 gravid females, 28 nongravid females, and 28 males) were collected from the vicinity of MLBS between 12 March 2025 and 28 March 2025. Females were candled to determine reproductive stage. Late stage eggs characteristic of gravid females in the spring are highly prominent due to their size (>1.5mm; gravid) and easily distinguished from absent or very early stage follicles (<1mm; nongravid) (Howard and Maerz 2022). Salamanders returning to the lab were processed as in Experiment 1 and all individuals had intact tails and (except for three non-gravid females) measured >34mm. However, we show that our main effects hold whether these three individuals (range = 32.53–33.50mm) are retained or eliminated from the dataset (see Appendix S2). Subjects were distributed evenly across the two thermal acclimation treatments with respect to size, sex, and reproductive class and acclimated for 19–29 days before starting physiological trials.

### iv. Stop-Flow Respirometry Protocol

Standard metabolic rates (SMR) were assayed as before except trials were only conducted at a single test temperature (16^°^C). This decision was made based on results of Experiment 1 suggesting that differences between groups are most pronounced at high test temperatures. Thus, each group of N=7 test subjects underwent a single day of trials and resumed normal feeding immediately afterwards. One subject (a non-gravid female in the warm thermal treatment) did not complete its trial due to mortality. All subjects were released at their point of capture at the conclusion of their trial.

### v. Statistical Analyses

To analyze variation in SMR across reproductive classes and treatments, we ran a series of linear mixed effects models as in Experiment 1, specifying log (SMR) as the response variable and trial date as a random effect. Mass-scaling coefficients were derived as before, and we screened for interaction effects before deciding to retain body mass as a covariate in all subsequent models. Differences in SMR between reproductive classes and treatments were visualized using mass-adjusted estimates, which were derived by dividing VO2 in ul/hr by starting body mass. To test for significant thermal acclimation responses to treatment and determine whether these differed between reproductive classes, we expanded our base model to include reproductive class, treatment, and their interaction as fixed effects. To identify significant pairwise differences in SMR across reproductive classes and treatments, post-hoc tests were carried out using the package emmeans (Lenth et al. 2022). Finally, to test for significant variation in SMR and thermal responses across the reproductive season, we combined all SMR estimates obtained from tests at 16^°^C and fitted separate models for each sex, which included season, treatment, and their interaction as fixed effects.

## Results

Subjects in the three reproductive classes differed significantly in mass (F = 26.468, *P* < 0.0001), with gravid females (1.195 ± 0.215 g) weighing significantly more than either non-gravid females (0.896 ± 0.210) or males (0.895 ± 0.107). Effects of body mass on SMR did not differ significantly among treatments or sexes (X^2^ =2.544, *P* = 0.280; Fig. S2). We detected no significant overall effect of reproductive class on SMR (X^2^ = 1.758, *P* = 0.415) but there was a significant overall treatment effect (X^2^ = 23.927, *P* < 0.0001) as well as a significant interaction between class and treatment (X^2^ = 11.628, *P* = 0.003; Fig. 3). Specifically, gravid females in the warm thermal treatment showed significant reductions in SMR compared to counterparts in the cool thermal treatment (gravid females: Tukey ratio = 4.887, *P* = 0.0001; non-gravid females: Tukey ratio = 3.819, *P* = 0.004; males: Tukey ratio = 3.270; *P* = 0.020) and non-gravid females in the warm thermal treatment (Tukey ratio = 3.070, *P* = 0.035).

**Figure 3.**
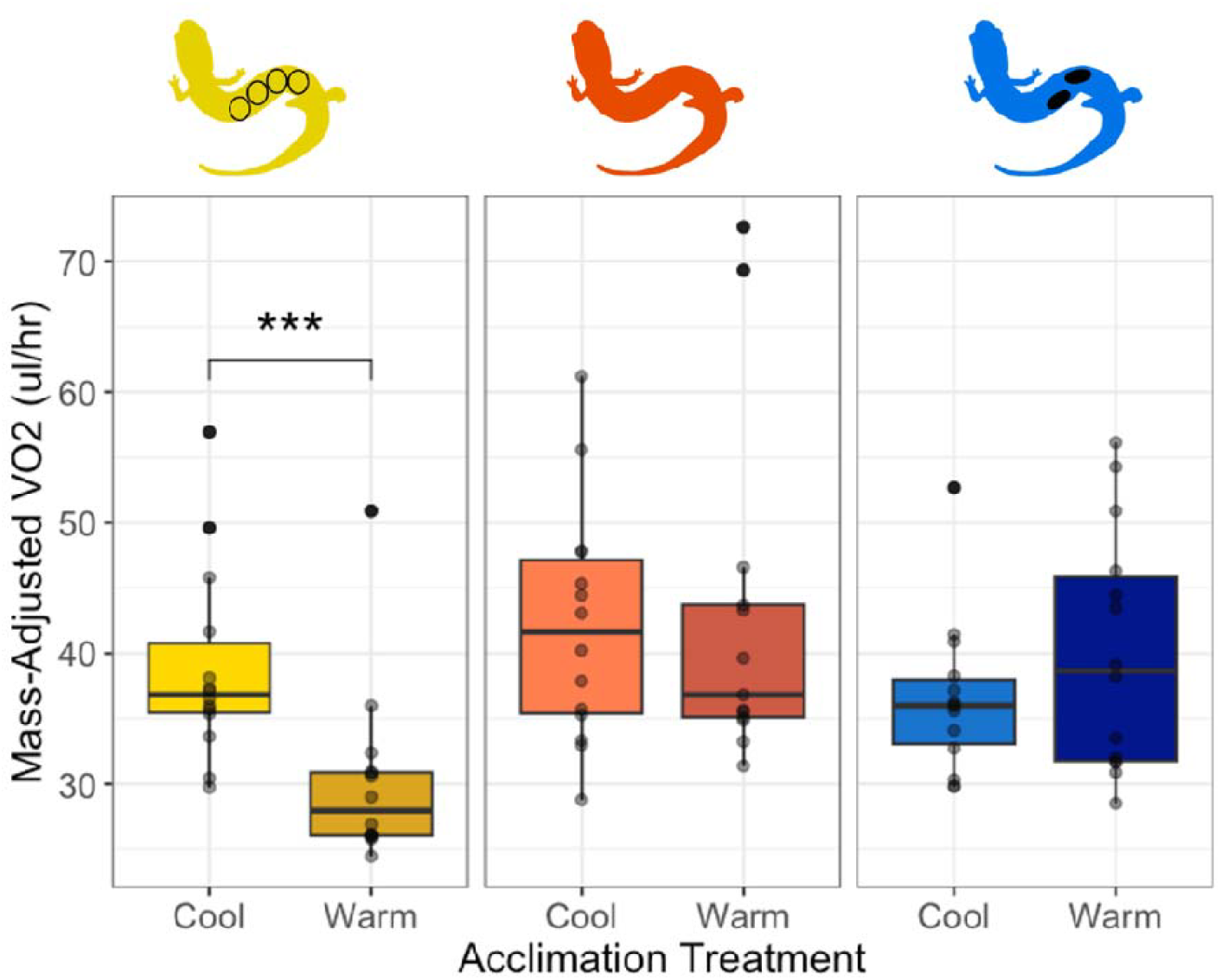
Standard metabolic rate (mass-adjusted VO2, in ul/hr) measured at 16^°^C in three classes of salamanders following three weeks of thermal acclimation at one of two treatments (Cool or Warm) in the spring: gravid females (N=14 cool, N=14 warm), non-gravid females (N=14 cool, N=13 warm), and males (N=14 cool, N=14 warm). Asterisks denote significant within-class treatment differences in a linear mixed effects model (* < 0.05, ** < 0.001, * * * < 0.0001).

### Seasonal Comparisons

To determine whether physiological rates differ across the breeding season, we compared SMR estimates at 16^°^C from gravid females and males assayed early (Fall) and late (Spring) in the breeding season (Fig. 4). Significant reductions in SMR were observed between the Fall and Spring in both gravid females (X^2^ = 5.486, P = 0.019) and males (X^2^ = 5.595, P = 0.018). We observed no significant interactions between season and treatment, suggesting that the pattern of response to thermal acclimation was consistent across seasons despite differences in base physiology.

**Figure 4.**
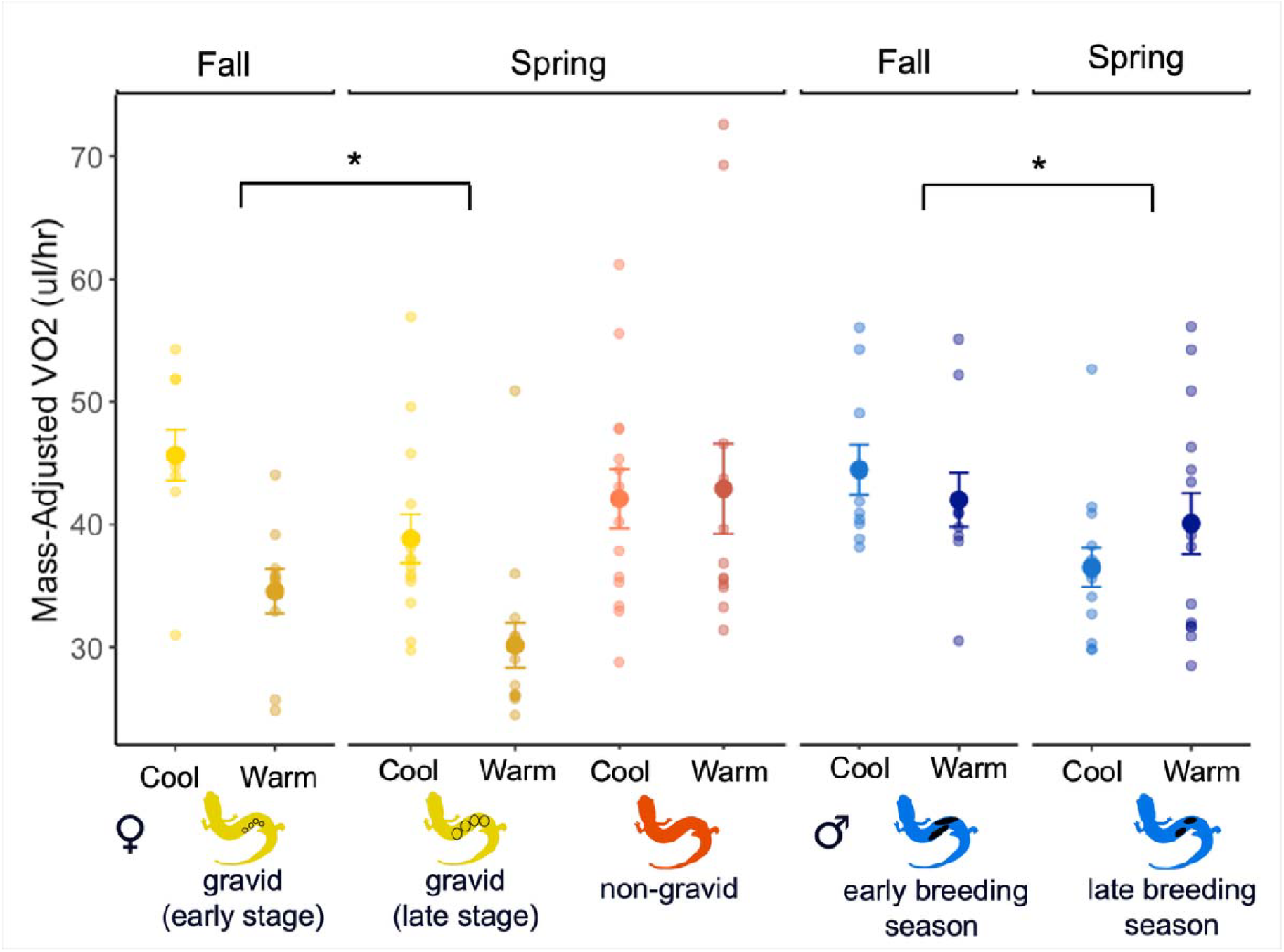
Standard metabolic rate (mass-adjusted VO2, in ul/hr) measured at 16^°^C at two points in the breeding season: Fall (September–November, early breeding season) and Spring (March–April, late breeding season). Three classes of salamanders are represented: Gravid females (in gold), non-gravid females (in coral), and males (in blue). Asterisks denote significant within-class season differences in a linear mixed effects model (* < 0.05, ** < 0.001, * * * < 0.0001).

## Discussion

The question of whether males and females differ in their responses to temperature change is essential to understanding adaptive capacities in a warming world, yet our basic understanding of when and where sex differences in thermal plasticity are likely to arise remains limited. Here, we tested for sex differences in thermal acclimation capacity in a plethodontid salamander, *P. cinereus*, across its protracted reproductive cycle. Our prediction was that reproductively active males and females would adopt distinct thermal acclimation strategies expected to optimize fitness relative to their sex-specific reproductive roles. We found patterns consistent with significant metabolic depression in gravid female, but not male salamanders, in response to warm temperature acclimation across the breeding season. Sex differences were not detected when sampling was restricted to non-gravid females at the same time of year, suggesting that patterns are driven by reproductive condition. Below, we discuss these results in the context of salamander thermal ecology and thermal ecology more broadly.

### No evidence for male-biased metabolic cold compensation during the breeding season

In cold-adapted ectotherms, enhanced physiological and/or behavioral plasticity of breeding males in response to low temperature acclimation has been suggested to convey sex-specific reproductive benefits, i.e., by enabling mating and courtship behaviors at temperatures that would otherwise be too cold to support activity (Rogers et al. 2007; Kiss et al. 2009; Moss et al. 2023). Given that the breeding season of P. cinereus at MLBS coincides with the fall and spring, when surface temperatures during nocturnal activity periods frequently drop below 5°C (Fig. 1A), we predicted that adult male salamanders acclimated to a cool thermal regime would show metabolic compensation at low temperatures compared to males acclimated to warmer temperatures. On the contrary, we found no evidence for metabolic compensation in males during the peak breeding season, with SMR at the coldest test temperature (8°C) being indistinguishable between thermal acclimation treatments. It is possible that sex differences in respiratory capacity at low temperatures manifest in more subtle ways than we examined here; for example, Rogers et al. (2007) observed no sex variation in whole-organism metabolism of frogs acclimated to autumn-like conditions but documented significant male bias in mitochondrial and enzyme activity in muscles used for calling and amplexus. Additional investigations may be warranted to investigate the sensitivity of specific, behaviorally relevant tissues to cold acclimation in salamanders.

### Female-biased metabolic depression at warm temperatures is associated with reproduction

Metabolic depression in response to sustained high temperature exposure is widespread among ectotherms, including lungless salamanders (Feder 1985; Bernardo and Spotila 2006; Markle and Kozak 2018; Muñoz et al. 2022), for which such plasticity is expected to enhance resilience by reducing maintenance energy costs and mitigating evaporative water loss (Riddell, Odom, et al. 2018; Riddell et al. 2019). In the current study, we observed metabolic depression in female, but not male P. cinereus, in response to warm temperature acclimation. Significant differences in SMR were observed between acclimation treatments at 16°C, the highest test temperature, but not at 8°C or 12°C, consistent with metabolic depression (Fig. 2). This result held for animals tested in both the fall and spring and appears to be driven by female reproductive condition, as no sex bias was detected when considering only non-gravid females. Hence, it is the process of vitellogenesis specifically, not other life history stages supporting reproduction, that shape the observed female-biased thermal acclimation strategy. This result is interesting in the light of what we know about female plethodontid reproductive cycles. At higher latitudes and elevations, where short growing seasons limit food intake and much of vitellogenesis occurs overwinter, females must attain a critical minimum body size before they may reproduce (Fitzpatrick 1973; Feder 1983; Crespi 2001; Leclair et al. 2008), and frequently skip years of reproduction to recoup depleted fat reserves (Gillette 2003; Leclair et al. 2008; Takahashi and Pauley 2010). The paradoxical outcome of this gradual accumulation strategy is that by the start of the breeding season, additional food intake is no longer necessary for females to proceed with vitellogenesis and yolk uptake; stored fat is simply shunted from growth to reproduction (Crespi 2001).

We propose that the thermal environment could interact with female reproductive strategies to reinforce an extreme energy conservation approach during vitellogenesis, stimulating metabolic depression to mitigate higher rates of energy turnover and preserve energy allocation to reproduction independent of food intake. Our data stand in contrast to the limited number of studies to date on metabolism in gravid female salamanders, which report significantly elevated mass-adjusted SMR when compared to males and non-gravid females (Fitzpatrick 1973; Finkler and Cullum 2002). Clearly, more studies are needed to determine the precise contribution of thermal environment, as well as other ecological factors, to female energetics across the protracted reproductive cycle.

### Both sexes exhibit seasonal metabolic rhythms

While we did not observe any changes in the general pattern of thermal acclimation of either males or gravid females across the reproductive cycle, comparison of measurements taken between fall and spring did reveal seasonal metabolic rhythms. Specifically, salamanders of both sexes showed significantly lower temperature-specific SMR when measured late in the reproductive cycle (spring) compared to early in the reproductive cycle (fall). To our knowledge, this pattern has not been previously documented in P. cinereus and could offer intriguing insights into seasonal patterns of energy allocation and activity. While we originally predicted that males at our study location would show higher metabolic rates in the fall compared to the spring to align energetic expenditure with seasonal peaks in breeding condition (i.e., circulating testosterone peaks in October at our study location (Church and Okazaki 2002)), the same pattern was not anticipated for females, who have been reported to emerge from brumation early in spring to maximize foraging time (Crespi 2001). A recent study investigating seasonal acclimatization of metabolic rates in spring-breeding spotted salamanders (*Ambystoma maculatum*) revealed a similar pattern of reduced SMR in the spring relative to the fall (Giacometti and Tattersall 2025), suggesting that seasonal rhythms may be attributed more to replenishing energy stores ahead of winter brumation than to breeding activities specifically. More research is needed to understand the adaptive significance of seasonal variation in resting metabolism.

### Conclusions and Future Directions

Our study provides new insights into the drivers of intraspecific variation in thermal traits in amphibians, a poorly understood topic (Cocciardi and Ohmer 2024). In the last decade, lungless salamanders have become models for the study of thermal ecology owing to their heightened physiological sensitivity (Sears et al. 2019; Riddell et al. 2024); however, a majority of this work has focused on characterizing macrogeographic variation within and between species (Riddell and Sears 2015; Markle and Kozak 2018; Muñoz et al. 2022). The contributions of sex or reproductive condition to seasonal patterns of thermal acclimation is virtually unstudied in this group. Given that the plethodontid reproductive cycle spans the majority of the year and is generally synonymous with the active season, forecasting the potential impacts of increasing thermal variability on wild salamander populations requires accounting for reproductive condition, especially given female demographic dominance in most populations. Whereas a previous study considering only adult males and juveniles reported an absence of physiological acclimation to warming conditions in two cold-adapted populations of *P. cinereus* (Muñoz et al. 2022), here we report pronounced physiological acclimation responses in gravid females from a montane population, suggesting that different demographic classes are likely to vary in their adaptive capacities. More studies are needed to evaluate the generalizability of this result across the extensive geographic range of *P. cinereus*, and to examine how patterns of thermal acclimation map onto other performance traits of interest, such as surface activity, reproductive success, and survival.

More broadly, our findings speak to the importance of accounting for sex in studies of thermal ecology. Not unlike the majority of scientific disciplines (Tannenbaum et al. 2019), the field of thermal ecology has a long-held tradition of ignoring sex as an explanatory variable (Huey and Pianka 2007; Lailvaux 2007; Bodensteiner et al. 2021; Edmands 2021; Gunderson 2021; Pottier et al. 2021; Hangartner et al. 2022). Despite a recent surge of interest in sex-specific thermal traits yielding several meta-analyses (Huey and Pianka 2007; Edmands 2021; Pottier et al. 2021; Hangartner et al. 2022), systematic patterns remain elusive, offering few insights into when and how sex may be important for designing studies. By taking a hypothesis-driven approach in an amphibian with sexually dimorphic reproductive life histories, we show that sex does account for differences in thermal acclimation strategies, but that reproductive stage matters. Future empirical studies must take both variables into account when testing for sex variation in thermal ecology, especially if population growth is limited by fecundity selection. differences in thermal acclimation capacity.

## Data Accessibility Statement

For the purposes of submission, all data transformation macros, extracted raw data, and analysis scripts have been uploaded along with the manuscript files for review. Upon acceptance, all the aforementioned data will be made publicly available in the Data Dryad repository.

## Acknowledgements

We would like to thank Dr. Vincent Farallo, Dr. Eric Riddell, and the technical support staff at Sable Systems International, in particular Jaco Klok and Barbara Joos, for their generous guidance and advice on the methodologies used for this study. This study was supported by Virginia Tech start-up funds to J.B.M.

## Ethical Approval

All procedures were approved by the Virginia Tech Institutional Animal Care and Use Committee (Protocol #23-250).

