## Appendix S1, Appendix S2, Fig. S1, Fig. S2 for "Sex and reproductive condition shape thermal acclimation strategy in a plethodontid salamander"

**Supplemental Material**

**Appendix S1: Detailed stop-flow respirometry protocol**

To initiate trials, subjects were guided into breathable, vinyl mesh sleeves held in shape by a rigid, cylindrical plastic frame. This platform served to elevate the animal, restrict movement, and minimize stress during trials. A moistened kimwipe was attached lengthwise to the bottom of each sleeve to provide a saturated local environment. Sleeves were then loaded into test chambers (60 mL clear acrylic tubes) which were sealed on each end with rubber stoppers and connected to an 8-channel flow multiplexer (Sable Systems RM-8)

The flow multiplexer and chambers were placed inside of a darkened environmental chamber programmed to the desired test temperature on the day prior. A sub-sampling pump (Sable Systems SS4) was used to pump source air from inside the incubator through the test chambers to the gas analyzer at a rate of 80 ml/min. Before reaching the multiplexer, the airstream passed through a soda lime scrubber to remove background CO2, followed by a water bubbler to saturate the airstream. An analog interface was used to automate the cycle of switching between chambers. To ensure measurable oxygen consumption, each test chamber was sealed for one hour before measuring its contents. Measurements occurred over three minutes with one minute baseline measurements taken between each chamber measurement. Air exiting chambers was passed to a gas analyzer (Sable Systems FMS) to analyze the volume of CO2 produced and O2 consumed. A magnesium perchlorate scrubber placed between the WVP and CO2 sensors was used to remove excess water vapor, and a soda lime scrubber placed between the CO2 and O2 sensors was used to remove CO2. Trials were monitored with overhead infrared security cameras and data were recorded live using Expedata software. We followed established manual bolus integration calculations to estimate VO2 (Lighton 2019). A total of five cycles was recorded for each trial, with the first two cycles discarded to account for acclimation time. Of the remaining three measures for each individual, we retained the lowest VO2 value as our estimate of SMR.

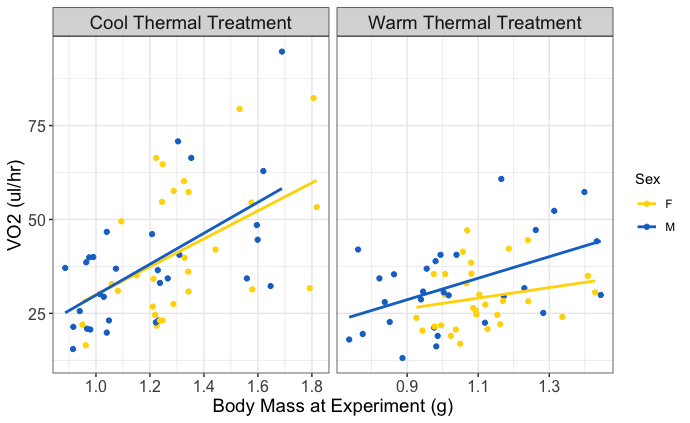

**Figure S1:** Body mass scaling relationships of SMR for Experiment 1. Mass-scaling coefficients were slightly elevated for warm-acclimated males (0.341) and cool-acclimated females (0.254) compared to cool-acclimated males (0.294) and warm-acclimated females (0.254; Fig. S1), although differences attributed to sex and treatment did not attain statistical significance (Χ^2^ = 0.008, *P* = 0.928).

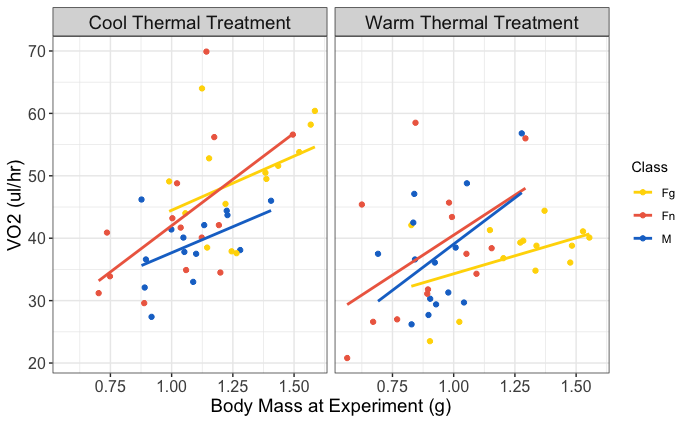

**Figure S2:** Body mass scaling relationships of SMR for Experiment 2. At 16°C, mass-scaling coefficients were steeper for non-gravid females (cool-acclimated = 0.350, warm-acclimated = 0.372) compared to males (cool-acclimated = 0.197, warm-acclimated = 0.251) or gravid females (cool-acclimated = 0.153 warm-acclimated = 0.165). However, effects of body mass on SMR did not differ significantly among treatments or sexes (Χ^2^ =2.544, *P* = 0.280)

**Main effect of Experiment 2 with n=3 nongravid females (measuring <34 mm) removed**

We detected no significant overall effect of reproductive class on SMR (Χ^2^ = 2.486, *P* = 0.289) but there was a significant overall treatment effect (Χ^2^ = 25.964, *P* < 0.0001) as well as a significant interaction between class and treatment (Χ^2^ = 13.801, *P* = 0.001). Specifically, gravid females in the warm thermal treatment showed significant reductions in SMR compared to counterparts in the cool thermal treatment (gravid females: Tukey ratio = 5.091, *P* = 0.0001; non-gravid females: Tukey ratio = 3.981, *P* = 0.002; males: Tukey ratio = 3.211; *P =* 0.024) and non-gravid females in the warm thermal treatment (Tukey ratio = 3.434, *P* = 0.0128).
